# Medial prefrontal decoupling from the default mode network benefits memory

**DOI:** 10.1101/706051

**Authors:** N.C.J. Müller, M. Dresler, G. Janzen, C.F. Beckmann, G Fernández, N. Kohn

## Abstract

In the last few years the involvement of the medial prefrontal cortex (mPFC) in memory processing has received increased attention. It is centrally involved when we use prior knowledge (schemas) to improve learning of new material. With the mPFC also being one of the core hubs of the default mode network (DMN) and the DMN’s role in memory retrieval, we decided to investigate whether the mPFC in a schema paradigm acts independently of the DMN. We tested this with data from a cross-sectional developmental study. During retrieval of schema items, the mPFC decoupled from the DMN with the degree of decoupling predicting memory performance. This finding suggests that a demand specific reconfiguration of the DMN supports schema memory. Additionally, we found that in the control condition, which relied on episodic memory, activity in the parahippocampal gyrus was positively related to memory performance. We interpret these results as a demand specific network reconfiguration of the DMN: a decoupling of the mPFC to support schema memory and a decoupling of the parahippocampal gyrus facilitating episodic memory. This supports the notion of dynamic reconfiguration of brain networks in response to task demands in the sense of process specific alliances.

## 1. Introduction

Declarative memory has most often been associated with activity and plasticity of the medial temporal lobe. However, in the last years, with the introduction of the ideas of schemas in cognitive neuroscience (Tse et al., 2007; van Kesteren et al., 2010a, 2012; Ghosh and Gilboa, 2014), the medial prefrontal cortex (mPFC) received more attention in the context of mnemonic processes. A schema is a knowledge structure that guides encoding, consolidation and retrieval of information that relates to prior knowledge (the schema) (Ghosh and Gilboa, 2014). The standard model of memory consolidation postulates that memories depend initially on the hippocampus and over time, in interaction with the prefrontal cortex, can be retrieved from neocortical representations independently of the hippocampus (Frankland and Bontempi, 2005; Takashima et al., 2006). Having a matching schema seems to dramatically accelerate the process of consolidation (Tse et al., 2007; van Kesteren et al., 2012). For this acceleration the mPFC is thought to link the new information to relevant schema (van Kesteren et al., 2010a; Tse et al., 2011). Thus, to understand the human memory better it will help to understand how the dynamics of the mPFC affect memory consolidation.

When investigating the mPFC in memory processing, there are at least two difficulties. First, the mPFC is a functionally heterogeneous area. It is not only implicated in schema processing, but also in other areas of cognition such as for example social cognition (Schilbach et al., 2008) and affective processing (Roy et al., 2012). Second, it is one of the core hubs of the default mode network (DMN), a network that deactivates when any task is performed and it activates during rest (Raichle et al., 2001; Raichle, 2015). Its activation during rest has been linked to self-referential thought (Gusnard et al., 2001), to mind-wandering (Mason et al., 2007) and autobiographic memory retrieval (Spreng and Grady, 2010). Especially the DMN’s involvement in mnemonic processes suggests a relation between the mPFC in schema memory and its function as a major hub of the DMN (Andrews-Hanna et al., 2010). Additionally, activity of the DMN under rest is linked to mind wandering (Mason et al., 2007). Mind wandering depends on recalling memories and constructing a narrative to “wander in”. Schema memory relies in a quite similar fashion on recalling information from the schema, which then is integrated with information that needs to be encoded or recalled. This analogy in processing makes it likely that the mPFC is not just coincidentally involved in the DMN and schema memory.

A second argument for an interrelation of the DMN and task activation of the mPFC is the strong coupling between resting state dynamics and task activation in general. We know for some time that on average there is a strong overlap between resting state networks and task dynamics (Smith et al., 2009). Modulation of DMN connectivity with other networks during goal-directed cognition (Spreng and Grady, 2010), after and during stress induction (Clemens et al., 2017; Young et al., 2017). Recently it has been shown that individual task activation can be accurately predicted from resting state data (Cole et al., 2016; Tavor et al., 2016). That predicting task activation is possible based on resting state alone suggests resting state functional connectivity forms the basis of task related activity. Therefore, taking into account functional connectivity reflected in resting state networks, might help us to get a better understanding of task related processing. One paper that nicely illustrates this is an investigation into the default mode network while performing a very simple finger opposition paradigm (Vatansever et al., 2015). The authors showed that the DMN reconfigured itself during the task, notably changing its connectivity towards the somatomotor network. During task performance (and fixation parts between trials) connectivity of the DMN to the left superior frontal gyrus correlated with behavioural performance suggesting a beneficial network reconfiguration. This study supports the notion that resting state networks reconfigure themselves in a demand specific fashion to support behaviour (Hasson et al., 2009). The idea of flexible reconfiguration of brain systems lies at the heart of the concept of process specific alliances; sets of brain regions that show elevated connectivity under task demands (Cabeza and Moscovitch, 2013). Recent evidence points to flexibility in DMN network connectivity in relation to episodic memory retrieval and future imaging (Bellana et al., 2017).

Recalling a schema memory activates the mPFC, a core hub of the DMN. Schema memory might only be a feature of the DMN, as schema might merely represent a self-referential knowledge systems that relates to previously established DMN functions. MPFC activity would then reflect a modulation of the DMN as a whole, with the mPFC being most visible as one of its core hubs. Alternatively, the mPFC, in an adaptive fashion in the sense of process specific alliances, decouples itself from the DMN to enable and facilitate schema memory. To investigate this, we re-analysed data from a large cross-sectional developmental study. In this study, participants first acquired a schema and then later learned new information. They could relate the information either to the schema (schema condition) or not (control condition). We tested whether the mPFC decouples itself from the default mode network under the schema condition relative to the control condition.

## 2. Methods

### 2.1 Participants

Ninety right-handed native Dutch-speaking volunteers participated in the original study, which tested three different age groups: Thirty adults aged between 25-32 years old (M = 26.9 years, SD = 21.9 months, 12 male), twenty-nine adolescents aged 18 (M = 18.5 years, SD = 3.1 months, 10 male) and thirty-one children aged between 10-12 years old (M = 11.0 years SD = 8.8 months, 8 male). All subjects had normal hearing and normal or corrected-to-normal vision. All participants were required to have no history of injury or disease known to affect the central nervous system function (including neuropsychological disorders such as dyslexia, autism and ADHD) and to not have MRI contraindications. Adults and adolescents were recruited from the student population of Radboud University, Nijmegen, and from the surrounding community. Children were recruited through presentations and flyers at local schools. The study was approved by the institutional Medical Research Ethics Committee (CMO Region, Arnhem-Nijmegen). Written informed consent was obtained prior to participation from all participants who were at least 18 years old; for the children participating both parents signed the informed consent. Of these 90 participants 3 children had to be excluded (1 did not want to complete the study, 1 moved excessively in the scanner, 1 due to an experimenter error); 2 adolescents were excluded as they did not complete the training at home; 1 adult had to be excluded due to an experimenter error. Of these 83 participants that completed the study we excluded 11 (6 children, 1 adolescent, 4 adults) participants that had fewer than 7 correct trials in one of the condition. This was done to prevent the high variability of fMRI activation in those participants to influence the results. All analysis focussed on this final set of 72 participants (21 children, 26 adolescents and 25 adults).

### 2.2 Behaviour

For this study we used data from a cross-sectional developmental study (see manuscript submitted in parallel). We decided to use this sample for two reasons. Not only was the task well suited, in the context of the study multiple measures were acquired. These additional measures could help to characterise any effect we observe better by relating it to behaviour. Furthermore, due to the study’s developmental dimension the sample size was substantially bigger compared to most studies. We summarise the design here as far as it is necessary to understand it for the analysis and results. The details can be found in another publication, which focused on the task analysis of this sample (Müller et al., 2019) or the initial publication of the task (van Buuren et al., 2014). The task was an object-location paradigm, similar to the game of “Memory/concentration“. Participants had to learn the layout of two boards containing in total 160 object-location associations. One board would contain the schema condition in which participants could use their prior knowledge to incorporate new associations in their schema, whereas the other board – the no-schema board – was a control condition. Over the first four days participants systematically learned the layout of the initial 40 schema associations; forming the experimental schema. On the no-schema board this systematic learning was prevented by the daily shuffling of the 40 control associations. In each session each object was presented three times followed by feedback about the correct location. On day five the 80 new associations were added on both boards: The 40 new schema, and the 40 new control associations. On the schema board participants could use their schemas to integrate the new associations whereas on the no-schema board this was not possible as the positions were randomly shuffled every day. Two days later participants returned to the lab and performed the retrieval test for all 160 associations. From the retrieval we obtained one score per condition: the schema associations, the control associations, the new schema and the new control associations. The trial structure during retrieval was as follows. One trial started with a cue, indicating the location for which object had to be retrieved, for 3s. Then, there was the response window of 3s followed by an inter trial interval with only a black fixation cross on screen for 2.5-7.5s. The inter trial interval was drawn from a uniform distribution. There was no feedback presented during recall. To keep the trial length and the visual input consistent across subjects the board would still be presented for the whole duration of the response window, independent of whether the response was already given. For relating behavioural performance to the decoupling of the mPFC we used two different aggregate scores: the schema benefit and a measure of general memory. The schema benefit quantifies how much participants profited from utilising the schema. It is calculated by subtracting the performance on the new associations in the control condition from that in the schema condition. As a measure of general memory performance independent of schema we calculated the average across the new associations. This score does not include the two training conditions as performance in those is not comparable due to the schema manipulation. Before calculating the average, we z-transformed both scores separately per group (children, adolescents and adults).

### 2.3 MRI data acquisition

Participants were scanned using a Siemens Magnetom Skyra 3 tesla MR scanner equipped with a 32-channel phased array head coil. The recall task comprised 935 volumes that were acquired using a T2* - weighted gradient-echo, multiecho echoplanar imaging sequence with the following parameters: TR = 2100ms; TE_1_ = 8.5ms, TE_2_ = 19.3ms, TE_3_ = 30ms, TE_4_ = 41ms; flip angle 90°; matrix size = 64 × 64; FOV = 224mm × 224mm × 119mm; voxel size = 3.5mm × 3.5mm × 3mm; slice thickness = 3mm; slice gap = 0.5mm; 34 slices acquired in ascending order. As this sequence did not provide whole brain coverage we oriented the FOV in a way that the hippocampus and the prefrontal cortex were fully inside and that only a small superior part of the parietal lobe was outside the FOV.

For the structural scans we used a T1-weighted magnetisation prepared, rapid acquisition, gradient echo sequence with the following parameters: TR = 2300ms; TE = 3.03ms; flip angle 8°; matrix size = 256 × 256; FOV= 192mm × 256mm × 256mm; slice thickness = 1mm; 192 sagittal slices.

### 2.4 MRI preprocessing

Preprocessing was done using a combination of FSL tools (Jenkinson et al., 2012), MATLAB (Natick, MA: The MathWorks) and ANTs (Avants et al., 2011a). From the two structural scans we generated an average using rigid body transformations from ANTs (Avants et al., 2011a), this procedure removed small movement induced noise. From the two scans and the average we always selected the scan with the least amount of ringing artefacts for all future analysis. If no difference was visible we used the average scan. These scans were denoised using N4 (Tustison et al., 2010) and generated a study specific template with an iterative procedure of diffeomorphic registrations (Avants and Gee, 2004). For the registration of the functional volumes we resampled the created template to a resolution of 3.5mm isotropic. Using Atropos (Avants et al., 2011b) the anatomical scans were segmented into 6 tissue classes: cerebrospinal fluid, white matter, cortical grey matter, subcortical grey matter, cerebellum and brainstem. The segmentation also produced individual brain masks.

For the functional multiecho data we combined echoes using in-house build MATLAB scripts. It used the 30 baseline volumes acquired during resting period directly before each part of the task to determine the optimal weighting of echo-times for each voxel (after applying a smoothing kernel of 3mm full-width at half-maximum to the baseline volumes), by calculating the contrast-to-noise ratio for each echo per scan. This script also directly realigned the volumes using rigid body transformation. Afterwards the volumes were smoothed using a 5mm full-width at half-maximum Gaussian kernel and grand mean intensity normalisation was done by multiplying the time series with a single factor. Younger participants tend to move more than older ones. For a developmental study it is thus important to minimise the effect of motion in the data. For this purpose we applied AROMA, a state of the art motion denoising algorithm that that uses independent component analysis decomposition of the data to identify movement and other noise signal (Pruim et al., 2015b, 2015a). Variance in the BOLD signal that could only be explained by components identified in this manner was regressed out. Afterwards we regressed out signal stemming from the cerebrospinal compartments and from the white matter by extracting the signal from individual generated segmentations using ANTs (Avants et al., 2011b). As a last step a 100s highpass filter was applied.

Boundary based registration was first calculated from native functional to native structural space using FLIRT (Greve and Fischl, 2009). We then calculated nonlinear registration from native structural space to the study template with FNIRT (Smith et al., 2004). The warping was done in a way that every functional volume was only resliced exactly once after the initial realignment. For displaying the final results we warped the final maps to MNI space using the nonlinear registration of ANTs (Avants and Gee, 2004).

### 2.4 MRI analysis

First, to establish whether the mPFC decouples itself from the DMN during the retrieval of schema items, we generated one mask for the DMN using the DMN mask from (Smith et al., 2009) and thresholded it at z>3.5. From that mask we then generated our second ROI, the mPFC mask, by removing all clusters except the one mPFC cluster. This enabled us to look at a specific decoupling of the anterior part of the DMN only. Then using simultaneous spatial regression with both masks as input, we generated one time series for each of the two ROIs. These time series reflect the dynamics of the mPFC corrected for the dynamics of the DMN and vice versa. We z-standardised both time-series by subtracting the grand mean and dividing by the standard deviation.

To compare the temporal dynamics between conditions and to generate peri-stimulus plots, we extracted a window of 7 volumes (12.6s in total) at the onset of each trial from each ROIs’ time series. These values were then averaged separately per condition.

To test the resulting curves for differences we used a repeated measures ANOVA with the 3 time points during which the hemodynamic response function should peak (4.2s, 6.3s, 8.4s). The model had the factors schema (schema & new schema vs. control & new control), initial or new (schema & control vs. new schema & new control), region (mPFC vs. DMN), time (4.2, 6.3, 8.4s) and to model age-related differences we included the between subject factor, group (children, adolescents, adults).

Additional whole brain analyses to validate localization of decoupling are described and presented in supplementary materials.

### 2.4 Assessment of behavioural relevance

In order to assess the behavioural relevance of this decoupling of the mPFC in the schema condition we tested whether the magnitude of decoupling in the schema vs. the control condition would predict memory performance.

### 2.4 Corollary analyses on memory systems underlying memory performance

As previous work has shown that the connectivity to the medial temporal lobe is gated through the parahippocampal gyrus (Ward et al., 2014), we tested whether it exhibits the complementary behaviour for the non-schema, general memory condition. Our mask was based on the Harvard-Oxford atlas included in FSL (Desikan et al., 2006).

## 3. Results

Comparing the dynamics of the mPFC and the DMN during the successful retrieval of object-location associations we found a condition specific dissociation of the mPFC when corrected for DMN dynamics. During retrieval of the schema associations, the activity level of the mPFC did not change whereas in the other conditions the mPFC deactivated with task onset; as the DMN did (Fig. 1). Focusing on the most relevant time window (trial onset + HRF delay) this effect is reflected in the three-way interaction of Region × Schema × initial/new (F(1,69)=16.1, p<.001, 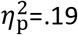). The three-way interaction is driven by the Schema × initial/new interaction in the mPFC (F(1,69)=60.57, p<.001, 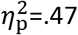) that is not evident in the DMN (F(1,69)=.06, p=.81, 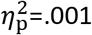). There was no indication of a significant difference of any of these effects between the different age groups. As we did not find any significant influence of age we collapsed across the three groups for the remaining analysis.

**Figure 1.**
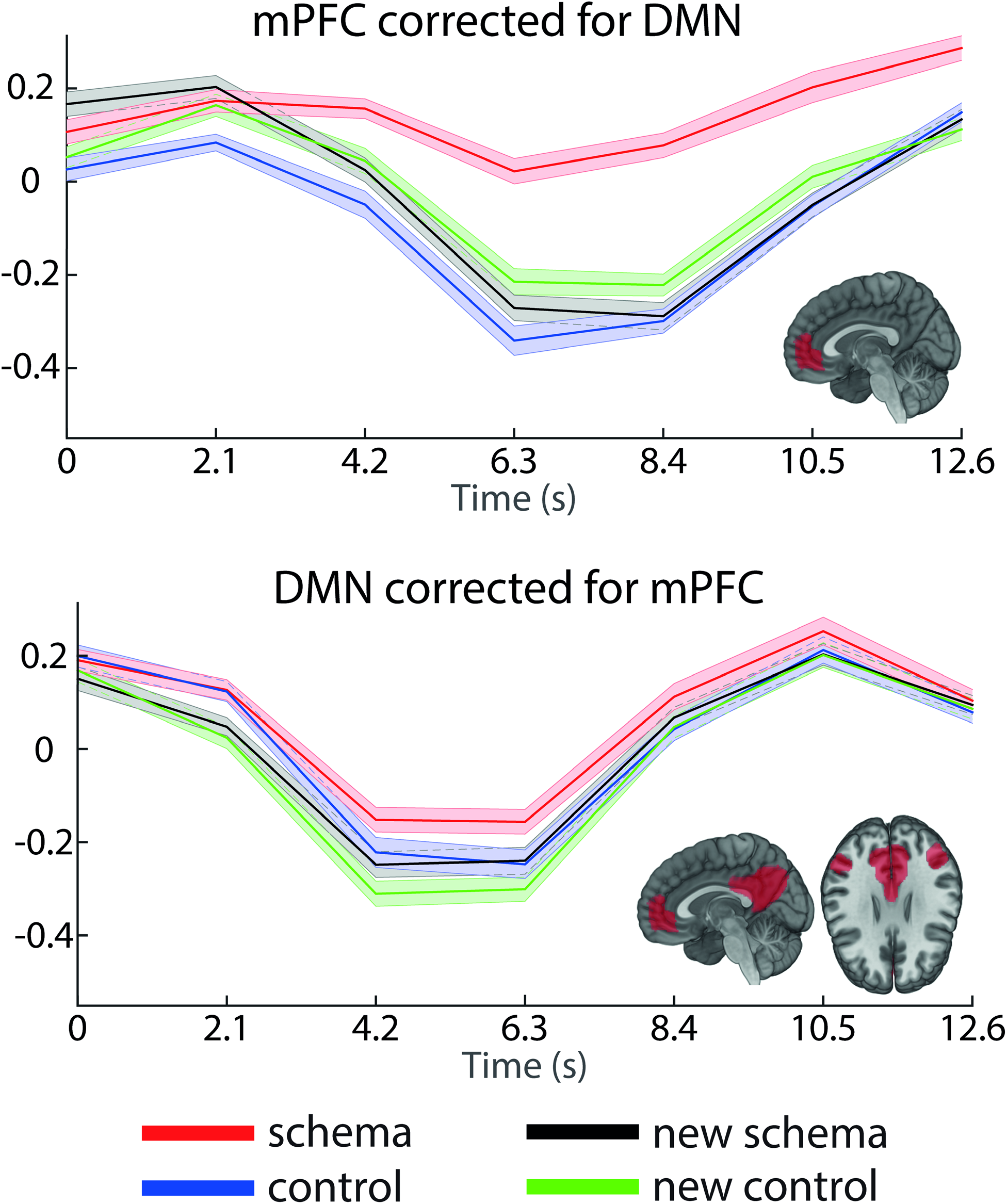
Decoupling of the mPFC during retrieval of schema memories. By simultaneously regressing the medial prefrontal cortex (mPFC) against the default mode network (DMN) time series and vice versa, we isolated their differential activations during the cued retrieval task. We found that across all conditions the DMN corrected for the mPFC shows its typical pattern of task-induced deactivation. The mPFC mostly followed that pattern, except for the condition of the schema associates in which its activation was not reduced as it was in the remainder of the DMN. The shaded area depicts the standard error of the mean. The y-axis depicts z-standardised activity.In assessing behavioural relevance, we indeed found positive correlations for the difference of the schema and control activation with the schema benefit (r(69)=.28, p=.02) and mean memory performance (r(69)=.24, p=.04); to be statistically more conservative, for both these correlations, we excluded one participant as his data would have increased the correlation substantially. Thus, recruiting the mPFC seems to have improved performance for the schema items. On top of that, participants that more adaptively recruited the mPFC in the schema over the control condition showed better memory overall.

In the behavioural assessment of memory systems underlying the two experimental conditions, we indeed found the exact opposite correlation: the difference control minus schema was positively correlated to the mean memory performance (r(69)=.29, p=.013, again the same participant excluded) in the parahippocampal gyrus. In conclusion, participants that activated the mPFC stronger in the schema over the control condition had better memory performance. Complementary, participants who activated the parahippocampal gyrus stronger in the control over the schema condition also showed better memory performance for the new paired associates.

## 4. Discussion

We found that the mPFC decoupled from the DMN during retrieval of schema memories. Whereas activity in the mPFC stayed constant during the schema condition, the posterior part of the DMN exhibited the typical task negative pattern. During retrieval of purely episodic memories in the control condition, both the mPFC and the posterior DMN deactivated synchronously. The stronger the mPFC decoupled in the schema relative to the control condition, the more a participant benefitted from schema and the higher was their overall memory performance. Additionally, stronger parahippocampal cortices decoupling was associated to better memory performance in the episodic memory related control condition. The decoupling of the mPFC and parahippocampal brain areas from the DMN suggests that schema facilitation is enabled by a specific reconfiguration of the DMN which is conceptually similar to process specific alliances (Cabeza and Moscovitch, 2013).

In previous research on the encoding or retrieval of schema memories, activation of the mPFC has repeatedly been associated with successful memory for schema items (van Kesteren et al., 2010b, 2010a, 2013, 2014; Liu et al., 2017). However, these papers did not report systematic activation of the DMN as a whole. Our account of the mPFC decoupling from the DMN during schemas are in line with a selective activation of the mPFC: If in those studies the mPFC decoupled from the DMN in the same way as we found, the mPFC would show a relative activation compared to a control condition whereas the remainder of the DMN would not show such activation. The decoupling of the mPFC could indicate a task-specific network reconfiguration of the DMN (Hasson et al., 2009; Cabeza and Moscovitch, 2013; Clemens et al., 2017). Vatansever and colleagues (Vatansever et al., 2015) demonstrated such a reconfiguration of the DMN during a finger opposition paradigm. During their task the posterior cingulate cortex, a hub of the DMN, changed its connectivity towards regions associated to motor planning. A dynamic re-coupling of the DMN during rest, retrieval of episodic memory and future thinking has previously been demonstrated (Bellana et al., 2017). We corroborate these findings and add the aspect of utility of this de-coupling in terms of benefit in memory performance. Analogously, two of the schema studies mentioned above indicate a reconfiguration of connectivity of the mPFC towards regions that encode the schema content. In one study, participants had to remember associations between textile fabric, visual patterns and the type of clothing they supposedly originate from (van Kesteren et al., 2010b): During the schema condition there was increased connectivity between the mPFC and a cluster in the somatosensory cortex, the stronger this connectivity the more the participants profited from the schema. In another schema study participants had to learn associations between famous faces and houses (Liu et al., 2017). For the schema condition, the study reported activation of the mPFC together with regions representing the stimulus content like the posterior place region and the fusiform face area. If the mPFC dynamic indeed reflects a task-specific reconfiguration, both these studies give an indication how this reconfiguration could look like: a shift away from the DMN network towards regions that contain the content of the schema. For example the tactile representation of the fabric (van Kesteren et al., 2010b) or the identity of the stimulus (Liu et al., 2017).

**Figure 2.**
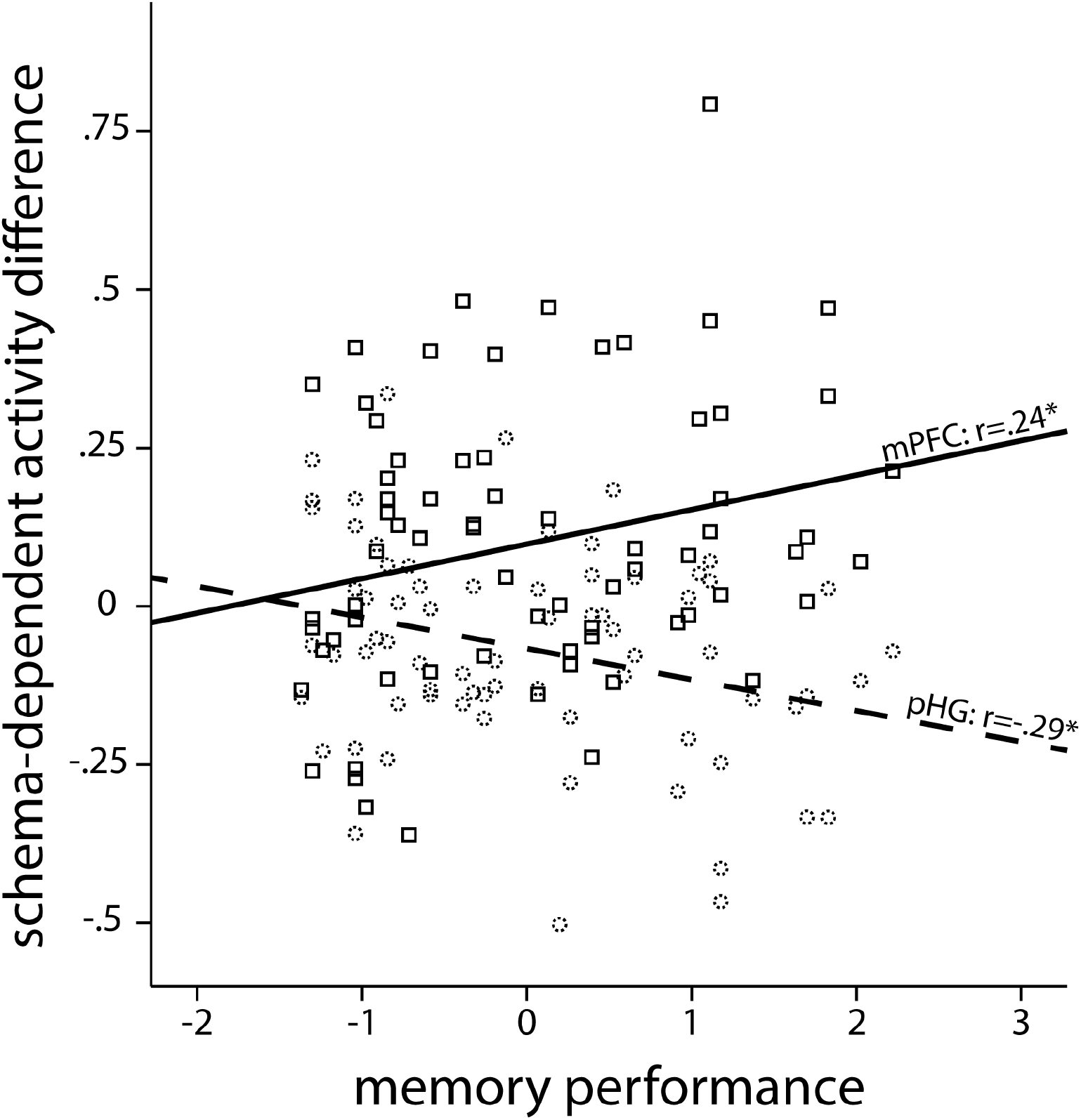
Decoupling of the mPFC during retrieval of schema memories. By simultaneously regressing the medial prefrontal cortex (mPFC) against the default mode network (DMN) time series and vice versa, we isolated their differential activations during the cued retrieval task. We found that across all conditions the DMN corrected for the mPFC shows its typical pattern of task-induced deactivation. The mPFC mostly followed that pattern, except for the condition of the schema associates in which its activation was not reduced as it was in the remainder of the DMN. The shaded area depicts the standard error of the mean. The y-axis depicts z-standardised activity.

When we found the localized decoupling of the mPFC, likely indicating an adaptive network configuration for schema processing, we wondered whether there would be a complementary reconfiguration of the DMN that benefited episodic processing during the no-schema condition. Therefore, we tested whether activity of the parahippocampal gyrus would also be associated to memory performance. We chose the parahippocampal gyrus as it is the primary hub of the DMN within the medial temporal lobe (Ward et al., 2014). Indeed, we found that its activity in the control condition over the schema condition positively predicted memory; the exact opposite relation compared to the mPFC. Bellana and colleagues (Bellana et al., 2017) similarly found that modulation of DMN coupling to medial temporal regions was associated to episodic memory retrieval.

The links between both the mPFC and the parahippocampal gyrus to memory performance were related to memory performance averaged across the new schema and the new control condition. We see this pattern of results as evidence for two different memory systems that are recruited in an adaptive fashion. With schemas available, the mPFC decouples whereas for episodic memories the parahippocampal gyrus decouples. The more strongly the mPFC and the parahippocampal gyrus decoupled, the more strongly the network reconfigured itself, the better the overall performance was. The results are in line with previously reported demand specific reconfiguration of the DMN being associated with good performance in a finger opposition (Vatansever et al., 2015) and a narrative comprehension task (Simony et al., 2016). Overall this shift, between a state that favours schema and one that favours episodic memory, suggests that schema processing is not an integral feature of episodic memory but rather a separate system that takes over if there is prior knowledge available as suggested previously (Fernández, 2017).

For our study, we used a developmental sample including children of around 10 years of age, adolescents that were 18 and adults that were above 25. Across these age ranges, we did not find any age related differences neither in the decoupling itself nor in the association of the decoupling to the performance. All groups showed the same dynamics. Only the children showed numerically smaller activity changes, those were not significant though. Of course, this null finding cannot be confidently interpreted in either direction, but it could potentially reflect that the DMN is by age 10 matured enough to enable this demand specific network reconfigurations (Fair et al., 2008). In particular, a previous longitudinal study did not find any maturational differences of the DMN in the mPFC cluster we found for the decoupling between the ages 10 and 13 (Sherman et al., 2014).

To conclude, we found that the retrieval of schema memory is not a feature of the DMN but is rather supported by a demand specific reconfiguration of the DMN. Such demand specific reconfigurations of large-scale brain networks have been called process specific alliances and were previously demonstrated for other memory and memory-related processes. During retrieval the mPFC decouples from the DMN to facilitate schema memory. Complementarily, in trials that relied on episodic memory, decoupling of the parahippocampal gyrus was associated with high levels of memory performance. Together, we interpret these findings as demand specific reconfiguration of the DMN. The better participants could adjust their network state based on task demands, the better they performed.

## 5. Acknowledgement

This research was funded by an NWO Research Talent Grant (406-13-008) to N.C.J.M. and G.F.

## References

Andrews-Hanna JR, Reidler JS, Sepulcre J, Poulin R, Buckner RL (2010) Functional-Anatomic Fractionation of the Brain’s Default Network. Neuron 65:550–562.

Avants BB, Gee JC (2004) Geodesic estimation for large deformation anatomical shape averaging and interpolation. Neuroimage 23:139–150.

Avants BB, Tustison NJ, Song G, Cook PA, Klein A, Gee JC (2011a) A reproducible evaluation of ANTs similarity metric performance in brain image registration. Neuroimage 54:2033–2044.

Avants BB, Tustison NJ, Wu J, Cook PA, Gee JC (2011b) An open source multivariate framework for N-tissue segmentation with evaluation on public data. Neuroinformatics 9:381–400.

Bellana B, Liu ZX, Diamond NB, Grady CL, Moscovitch M (2017) Similarities and differences in the default mode network across rest, retrieval, and future imagining. Hum Brain Mapp.

Cabeza R, Moscovitch M (2013) Memory Systems, Processing Modes, and Components: Functional Neuroimaging Evidence. Perspect Psychol Sci.

Clemens B, Wagels L, Bauchmüller M, Bergs R, Habel U, Kohn N (2017) Alerted default mode: functional connectivity changes in the aftermath of social stress. Sci Rep 7:40180.

Cole MW, Ito T, Bassett DS, Schultz DH (2016) Activity flow over resting-state networks shapes cognitive task activations. Nat Neurosci 19:1718–1726.

Desikan RS, Ségonne F, Fischl B, Quinn BT, Dickerson BC, Blacker D, Buckner RL, Dale AM, Maguire RP, Hyman BT, Albert MS, Killiany RJ (2006) An automated labeling system for subdividing the human cerebral cortex on MRI scans into gyral based regions of interest. Neuroimage 31:968–980.

Fair D a, Cohen AL, Dosenbach NUF, Church J a, Miezin FM, Barch DM, Raichle ME, Petersen SE, Schlaggar BL (2008) The maturing architecture of the brain’s default network. Proc Natl Acad Sci 105:4028–4032.

Fernández G (2017) The Medial Prefrontal Cortex is a Critical Hub in the Declarative Memory System. In: Cognitive Neuroscience of Memory Consolidation, pp 45–56. Springer International Publishing.

Frankland PW, Bontempi B (2005) The organization of recent and remote memories. Nat Rev Neurosci 6:119–130.

Ghosh VE, Gilboa A (2014) What is a memory schema? A historical perspective on current neuroscience literature. Neuropsychologia 53:104–114.

Greve DN, Fischl B (2009) Accurate and robust brain image alignment using boundary-based registration. Neuroimage 48:63–72.

Gusnard DA, Akbudak E, Shulman GL, Raichle ME (2001) Medial prefrontal cortex and self-referential mental activity: Relation to a default mode of brain function. Proc Natl Acad Sci 98:4259–4264.

Hasson U, Nusbaum HC, Small SL (2009) Task-dependent organization of brain regions active during rest. Proc Natl Acad Sci 106:10841–10846.

Jenkinson M, Beckmann CF, Behrens TEJ, Woolrich MW, Smith SM (2012) Fsl. Neuroimage 62:782–790.

Liu ZX, Grady C, Moscovitch M (2017) Effects of Prior-Knowledge on Brain Activation and Connectivity During Associative Memory Encoding. Cereb Cortex 27:1991–2009.

Mason MF, Norton MI, Horn JD Van, Wegner DM, Grafton ST, Macrae CN, Mason MF, Norton MI, Horn JD Van, Wegner DM, Grafton ST, Macrae CN (2007) Wandering minds: Stimulus-independent thought. Science 315:393–395.

Müller NCJ, Kohn N, van Buuren M, Klijn N, Emmen H, Berkers RMWJ, Dresler M, Janzen G, Fernández G (2019) Differences in strategic abilities but not associative processes explain memory development. bioRxiv:693895

Pruim RHR, Mennes M, Buitelaar JK, Beckmann CF (2015a) Evaluation of ICA-AROMA and alternative strategies for motion artifact removal in resting state fMRI. Neuroimage 112:278–287.

Pruim RHR, Mennes M, van Rooij D, Llera A, Buitelaar JK, Beckmann CF (2015b) ICA-AROMA: A robust ICA-based strategy for removing motion artifacts from fMRI data. Neuroimage 112:267–2774.

Raichle ME (2015) The Brain’s Default Mode Network. Annu Rev Neurosci 38:433–447.

Raichle ME, MacLeod AM, Snyder AZ, Powers WJ, Gusnard DA, Shulman GL (2001) A default mode of brain function. Proc Natl Acad Sci 98:676–682.

Roy M, Shohamy D, Wager TD (2012) Ventromedial prefrontal-subcortical systems and the generation of affective meaning. Trends Cogn Sci 16:147–156.

Schilbach L, Eickhoff SB, Rotarska-Jagiela A, Fink GR, Vogeley K (2008) Minds at rest? Social cognition as the default mode of cognizing and its putative relationship to the “default system” of the brain. Conscious Cogn 17:457–467.

Sherman LE, Rudie JD, Pfeifer JH, Masten CL, McNealy K, Dapretto M (2014) Development of the Default Mode and Central Executive Networks across early adolescence: A longitudinal study. Dev Cogn Neurosci 10:148–159.

Simony E, Honey CJ, Chen J, Lositsky O, Yeshurun Y, Wiesel A, Hasson U (2016) Dynamic reconfiguration of the default mode network during narrative comprehension. Nat Commun 7:12141.

Smith SM, Fox PT, Miller KL, Glahn DC, Fox PM, Mackay CE, Filippini N, Watkins KE, Toro R, Laird AR, Beckmann CF (2009) Correspondence of the brain’s functional architecture during activation and rest. Proc Natl Acad Sci 106:13040–13045.

Smith SM, Jenkinson M, Woolrich MW, Beckmann CF, Behrens TEJ, Johansen-Berg H, Bannister PR, De Luca M, Drobnjak I, Flitney DE, Niazy RK, Saunders J, Vickers J, Zhang Y, De Stefano N, Brady JM, Matthews PM (2004) Advances in functional and structural MR image analysis and implementation as FSL. Neuroimage 23:S208–S219.

Spreng RN, Grady CL (2010) Patterns of Brain Activity Supporting Autobiographical Memory, Prospection, and Theory of Mind, and Their Relationship to the Default Mode Network. J Cogn Neurosci 22:1112–1123.

Takashima a, Petersson KM, Rutters F, Tendolkar I, Jensen O, Zwarts MJ, McNaughton BL, Fernández G (2006) Declarative memory consolidation in humans: a prospective functional magnetic resonance imaging study. Proc Natl Acad Sci 103:756–761.

Tavor I, Jones OP, Mars RB, Smith SM, Behrens TE, Jbabdi S (2016) Task-free MRI predicts individual differences in brain activity during task performance. 352:1773–1776.

Tse D, Langston RF, Kakeyama M, Bethus I, Spooner PA, Wood ER, Witter MP, Morris RGM (2007) Schemas and memory consolidation. Science 316:76–82.

Tse D, Takeuchi T, Kakeyama M, Kajii Y, Okuno H, Tohyama C, Bito H, Morris RGM (2011) Schema-dependent gene activation and memory encoding in neocortex. Science 333:891–895.

Tustison NJ, Avants BB, Cook P a., Zheng Y, Egan A, Yushkevich P a., Gee JC (2010) N4ITK: Improved N3 bias correction. IEEE Trans Med Imaging 29:1310–1320.

van Buuren M, Kroes MCW, Wagner IC, Genzel L, Morris RGM, Fernandez G (2014) Initial Investigation of the Effects of an Experimentally Learned Schema on Spatial Associative Memory in Humans. J Neurosci 34:16662–16670.

van Kesteren MTR, Beul SF, Takashima A, Henson RN, Ruiter DJ, Fernández G (2013) Differential roles for medial prefrontal and medial temporal cortices in schema-dependent encoding: From congruent to incongruent. Neuropsychologia 51:2352–2359.

van Kesteren MTR, Fernández G, Norris DG, Hermans EJ, Fernandez G, Norris DG, Hermans EJ (2010a) Persistent schema-dependent hippocampal-neocortical connectivity during memory encoding and postencoding rest in humans. Proc Natl Acad Sci 107:7550–7555.

van Kesteren MTR, Rijpkema M, Ruiter DJ, Fernandez G, Fernández G (2010b) Retrieval of associative information congruent with prior knowledge is related to increased medial prefrontal activity and connectivity. J Neurosci 30:15888–15894.

van Kesteren MTR, Rijpkema M, Ruiter DJ, Morris RGM, Fernández G, Kesteren MTR Van, Rijpkema M, Ruiter DJ, Morris RGM, Fernández G (2014) Building on Prior Knowledge: Schema-dependent Encoding Processes Relate to Academic Performance. J Cogn Neurosci 26:2250–2261.

van Kesteren MTR, Ruiter DJ, Fernández G, Henson RN (2012) How schema and novelty augment memory formation. Trends Neurosci 35:211–219.

Vatansever D, Menon DK, Manktelow AE, Sahakian BJ, Stamatakis EA (2015) Default mode network connectivity during task execution. Neuroimage 122:96–104.

Ward AM, Schultz AP, Huijbers W, Van Dijk KR a, Hedden T, Sperling R a (2014) The parahippocampal gyrus links the default-mode cortical network with the medial temporal lobe memory system. Hum Brain Mapp 35:1061–1073.

Young CB, Raz G, Everaerd D, Beckmann CF, Tendolkar I, Hendler T, Fernández G, Hermans EJ (2017) Dynamic Shifts in Large-Scale Brain Network Balance As a Function of Arousal. J Neurosci 37:281–290.

